# Selection for insecticide resistance can promote *Plasmodium falciparum* infection in *Anopheles*

**DOI:** 10.1101/2022.12.07.516767

**Authors:** Kelsey L. Adams, Emily K. Selland, Bailey C. Willett, John W. Carew, Charles Vidoudez, Naresh Singh, Flaminia Catteruccia

**Author notes:** Corresponding author: Flaminia Catteruccia **Email:**. **Author Contributions:** K.L.A. contributed to literature searches, study design, data collection, data analysis, data interpretation, figure creation, and writing; E.K.S., B.C.W, J.W.C. and C.V. contributed to data analysis and data collection, N.S. contributed to data collection, and F.C. contributed to study design, data analysis, data interpretation, writing, and project supervision. **Competing Interest Statement:** The authors have no competing interests.

## Abstract

Insecticide resistance is under strong selective pressure in *Anopheles* mosquitoes due to widespread usage of insecticides in vector control strategies. Resistance mechanisms likely cause changes that profoundly affect mosquito physiology, yet it remains poorly understood how selective pressures imposed by insecticides may alter the ability of the mosquito to host and transmit a *Plasmodium* infection. From pyrethroid-resistant field-derived *Anopheles gambiae s*.*l*. mosquitoes, we performed selection experiments to establish resistant (RES) and susceptible (SUS) colonies by either selection for, or loss of, insecticide resistance. We show increased prevalence, intensity, and oocyst growth rate of *Plasmodium falciparum* infection in RES females compared to SUS. The increase in infection intensity in RES females was not associated with the presence of the *kdr*L1014F mutation, and was not impacted by inhibition of Cytochrome P450s. The lipid transporter lipophorin (Lp), which was upregulated in RES compared to SUS, was at least partly implicated in the increased intensity of *P. falciparum* but not directly in the insecticide resistance phenotype. Interestingly, we observed that although *P. falciparum* infections were not affected when RES females were exposed to permethrin, these females had decreased lipid abundance in the fat body following exposure, pointing to a possible role for lipid mobilization in response to damage caused by insecticide challenge. The finding that selection for insecticide resistance can increase *P. falciparum* infection intensities and growth rate reinforces the need to assess the overall impact on malaria transmission dynamics of selective pressures mosquitoes experience during repeated insecticide challenge.

**Significance Statement:** Insecticide resistance poses a severe threat for malaria control. Resistance to pyrethroid insecticides, the active component of most insecticide-treated nets, is now widespread in sub-Saharan Africa, reducing the efficacy of these crucial tools. Despite significant research characterizing insecticide resistance mechanisms, it remains unknown how these traits influence *Plasmodium falciparum* infections in malaria-transmitting *Anopheles* mosquitoes. We established a pyrethroid-resistant and pyrethroid-susceptible population of *Anopheles gambiae* derived from the same genetic background and performed experimental infections with *P. falciparum*. We found that the pyrethroid-resistant population was more supportive of malaria parasites compared to the susceptible population. This was not caused by well-known insecticide resistance mechanisms, but linked with a lipid transporter, lipophorin, which may play an indirect role in resistance.

## Introduction

Insecticide resistance is widespread in *Anopheles* populations and represents a major threat to vector control in sub-Saharan Africa and other malaria-endemic regions ^1^. The greatest reductions in the malaria burden in the last 20 years have been facilitated by long-lasting insecticide-treated nets (LLINs) impregnated with pyrethroids ^2^. Pyrethroids were the only class of insecticides approved for use in these tools until recently and continue to be the most dominant ^1-3^. Such limited chemical diversity has promoted insecticide resistance in *Anopheles* populations, to the point where there are now sparse reports of pyrethroid susceptibility in sub-Saharan Africa (irmapper.com). Not only is the prevalence of insecticide resistance increasing, but in some regions, vectors are able to withstand extremely high doses of pyrethroids. For example, in Burkina Faso, an *Anopheles coluzzii* population was demonstrated to increase its resistance intensity by 2.6-fold over a single malaria transmission season, and could withstand >1000X the lethal insecticide dose for a susceptible population ^4^.

Despite its widespread prevalence, many aspects of insecticide resistance remain poorly understood in mosquitoes. Some resistance mechanisms have been only recently described ^5-7^ while some others were long recognized ^8-13^, but their relative contribution to resistance and their overall physiological impacts remain to be deciphered. The most commonly reported mechanism, due to its ease of detection ^10,11^, is target site resistance, which relies on amino acid substitutions in the molecular target of the insecticide (knock down resistance *(kdr)* mutations in the voltage-gated sodium channel *para* for pyrethroids and organochlorines, and acetylcholinesterase mutations (*ace-1*) for carbamates and organophosphates) ^8,9,14^. Metabolic resistance is also well described, involving upregulated transcription and/or boosted activity of enzymes such as cytochrome P450s (CYPs) or glutathione-S-transferases (GSTs) that contribute to xenobiotic detoxification ^12,13,15^. Less well studied, cuticular resistance is characterized by a thickened cuticle caused by increased deposition of cuticular hydrocarbons (CHCs) or proteins that limit insecticide penetration ^7,16^.

We have only a narrow understanding of how these physiological adaptations affect other biological processes in the mosquito. Introgression of *kdr* mutations in the voltage-gated sodium channel has been associated with decreased fecundity, reduced mating success, and slower larval development in *Aedes aegypti* ^17^, and similar costs were seen associated with a metabolic resistance gene in *Anopheles funestus* ^18,19^ and in a resistant strain of *Aedes albopictus* ^20^. Impacts of insecticide resistance mechanisms on infections with the human malaria parasite *Plasmodium falciparum* are also not well understood, as reviewed extensively elsewhere ^21^. In *Anopheles gambiae*, introgression of the *kdrL1014F* target site resistance allele into a susceptible background caused a decrease in fecundity and also presented interactive costs between *P. falciparum* infections and *kdr*, showing a decrease in longevity of unexposed mosquitoes carrying these mutations but only when females were infected ^22^. In terms of a direct impact on infection outcomes, some observations have also associated *kdr*L1014F in *An. gambiae* with higher *P. falciparum* prevalence or intensity ^23-25^, while in another study no relationship was observed ^26^. Similar conflicting evidence surrounds a mutation in the metabolic resistance enzyme *GSTe2* in wild *An. funestus*, where the same polymorphism was associated with reduced oocyst prevalence ^27^ and higher sporozoite prevalence ^28^ in two different studies. There is also some evidence that sublethal insecticide exposure can decrease *P. falciparum* burdens ^*29-31*^ but it is not understood how these effects are mediated.

The lack of knowledge regarding how most metabolic and cuticular resistance mechanisms impact vector competence of *Anopheles* mosquitoes is particularly surprising considering these mechanisms are intertwined with mosquito processes that influence *Plasmodium* infection. Indeed, CYPs and GSTs catalyze enzymatic reactions that can both produce superoxide radicals (in the case of many CYPs) ^32^ or act as reducing agents to counteract oxidative stress (in the case of GSTs) ^33,34^, overall altering the redox state, which, in the midgut, regulates *Plasmodium* infections ^32-36^. Decreased energy reserves were observed in *Culex pipiens* mosquitoes that overexpressed CYPs, and it is also possible that resource availability and trade-offs could impact pathogen development ^37,38^. In cuticular resistance, CHC production utilizes lipid biosynthesis enzymes and lipid transporters ^39-41^ that are thought to be exploited by parasites ^42-44^, with potential for resource allocation to have consequences for *Plasmodium* infections ^42-44^.

The multi-locus nature of these resistance mechanisms creates a challenge for clearly addressing these questions. We approached this challenge by developing permethrin resistant (RES) and susceptible (SUS) colonies derived from the same genetic background: a highly resistant field population of *An. coluzzii* in Burkina Faso which was outcrossed to a susceptible lab population prior to selection (or deselection) for pyrethroid resistance. We determined that RES females possess both metabolic (CYPs) and target site (*kdr*) resistance, and support faster oocyst development, increased oocyst intensities, and higher prevalence and intensity of sporozoites. These effects on parasite survival and development appear to be independent of known target site and metabolic insecticide resistance mechanisms, but are associated with the function of the lipid transporter Lipophorin (Lp). Permethrin exposure in RES did not impact infection but led to decreased lipid levels in the fat body, potentially implicating lipid mobilization by Lp in the response to damage caused by insecticide challenge. Our results reveal that selection for permethrin resistance can – likely indirectly in this case– promote *Plasmodium* infection, an important finding at a time when insecticide pressure in field populations is nearly ubiquitous.

## Results

### Generating resistant and susceptible lines in a comparable genetic background

To investigate how insecticide resistance influences *Plasmodium* infections, we set out to compare two *Anopheles* lines that share a similar genetic background but differ in their phenotypic display of resistance to pyrethroids. We generated these colonies from a field-derived, highly pyrethroid-resistant parental *An. coluzzii* strain originating from the Vallée du Kou region VK5 in Burkina Faso **(Figure 1A, B, Supplementary Figure 1A)** recently adapted to lab conditions (VK strain), which demonstrated comparable resistance intensities to the original field mosquitoes **(Figure 1C)**. We could not deselect insecticide resistance in VK even after 38 generations **(Supplementary Figure 1A, B)**, so we decided to outcross this strain to a fully susceptible colony of *An. gambiae* (G3) and perform selection experiments in the ensuing strain (G3VK) **(Figure 1A)**. To this aim, we selected one branch with permethrin exposure every generation (RES) and removed insecticide pressure from the other (SUS). RES showed increasing resistance intensity using 1X (0.75%) and 5X (3.75%) doses of permethrin in the WHO bioassay, and reached resistance levels similar to its parental VK colony (after 18 generations), at which point the SUS colony showed complete loss of resistance **(Figure 1D)**.

**Figure 1:**
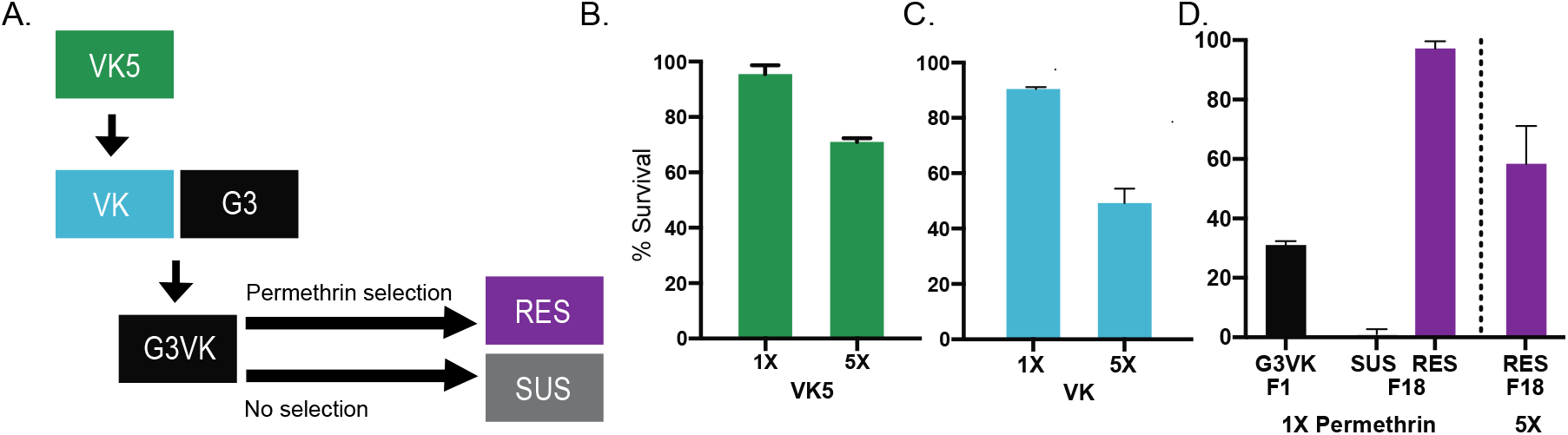
Generating resistant and susceptible colonies with a similar genetic background. (A) Schematic of crosses to generate resistant (RES) and susceptible (SUS) lines of *An. gambiae s*.*l*. VK was colonized from wild VK5 females in Burkina Faso. VK was outcrossed to G3 to create G3VK which was used in selection experiments generating RES and SUS. Survival was measured 24h post exposure to either a 0.75% (1X) and 3.75% (5X) for 1h in (B) parental wild VK5 females, (C) the VK-derived lab colony, and (D) the original G3VK F1 progeny of VK and G3 parents, and the subsequent RES and SUS colonies derived from the selection.

### Selected RES mosquitoes show multiple mechanisms of insecticide resistance

After selection, we ascertained which mechanisms of insecticide resistance were at play in RES, investigating a possible role for target site, metabolic and cuticular resistance. We determined that the common target site resistance mutation *kdrL1014F* is present in the RES colony at an allele frequency of 52% while it is nearly absent from the SUS colony **(Figure 2A)**. We only investigated *kdr*L1014F, as it is most dominant in VK5 field populations ^47^, but other mutations in *para* have also been described to a lesser extent in nearby regions ^48^. To assess the role of metabolic resistance, we first investigated whether GSTs or CYPs contribute to survival upon insecticide exposure in RES females. We found that, while pre-treatment with the GST inhibitor diethyl maleate (DEM, 8%) had no effect on susceptibility, pre-treatment with piperonyl butoxide (PBO, 4%) increased mortality of RES mosquitoes during permethrin exposure **(Figure 2B, C)**, implicating increased CYP activity in the resistance phenotype. Consistently, transcriptional analysis confirmed that *CYP6P3* and *CYP6Z2*, CYPs commonly associated with pyrethroid resistance ^12,45,46^, were both upregulated in RES, while we did not detect differences in *CYP6M2* **(Figure 2D)**. This is also the case in the parental VK5 population ^47^, providing a reassuring indication that relevant resistance mechanisms from the field have been conserved through the selection process. Finally, we did not see upregulation of *CYP4G16* and *CYP4G17*, two canonical markers of cuticular resistance that act as decarbonylases in the CHC biosynthesis pathway^6,7^ (**Figure 2E)**, suggesting that cuticular resistance via increased CHCs production does not mediate resistance in the RES colony.

**Figure 2:**
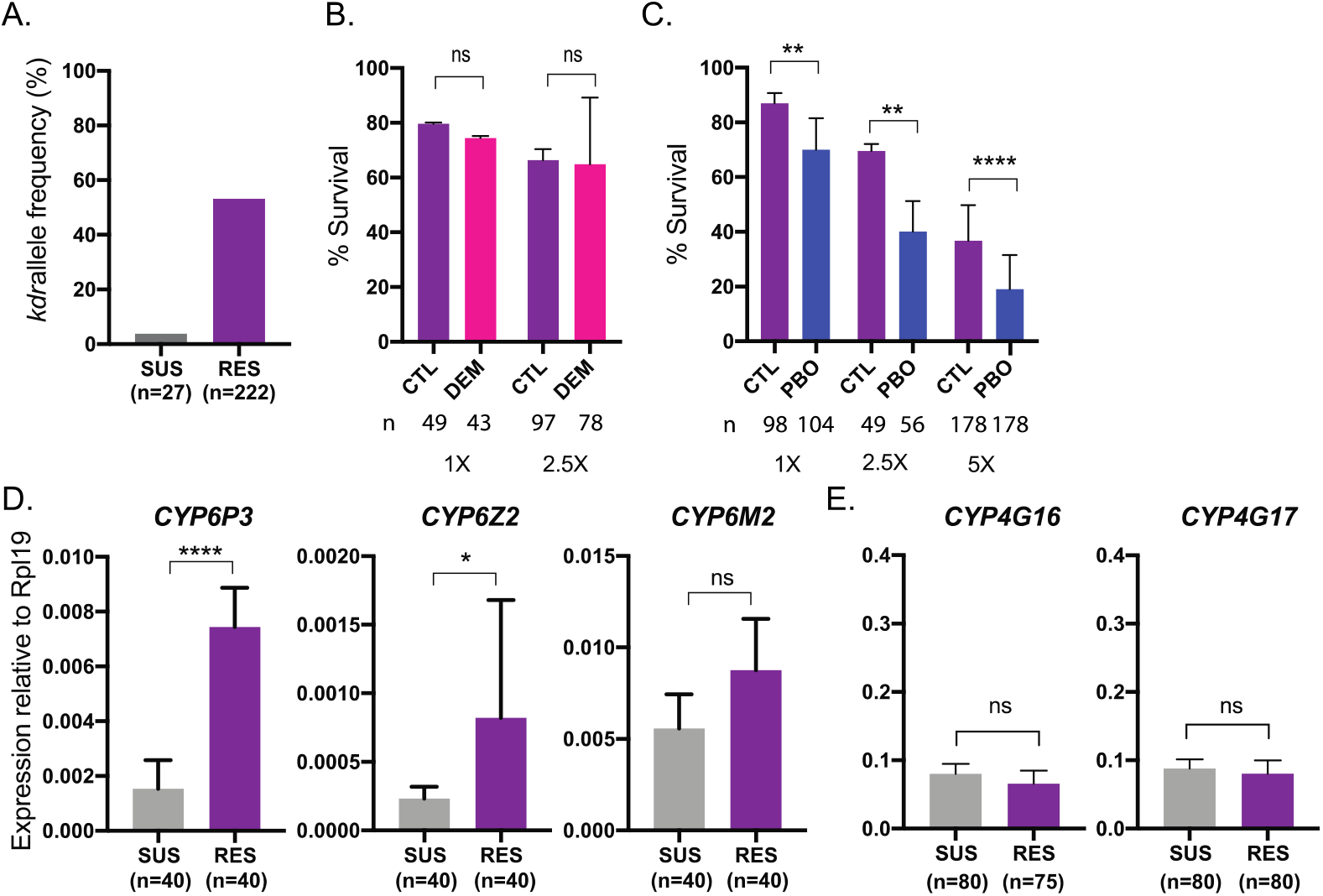
RES females have target site and metabolic resistance mechanisms. (A) The *kdrL1014F* mutation in the RES colony is at an allele frequency of 52% while only at 3% in the SUS colony. (B) Pre-exposure to 8% DEM did not impact mortality of females when exposed to either a 1X or 2.5X dose of permethrin (Chi-squared tests, *p*>0.05). Mean and SD are shown. (C) Pre-exposure to 4% PBO increased mortality of RES females when exposed to permethrin at either a 1X, 2.5X or 5X dose (Chi-squared tests, *p*=0.004, *p*=0.002, *p*<0.0001, respectively). Mean and SD are shown. (D) Transcript levels of metabolic resistance enzymes *CYP6P3* and *CYP6Z2*, but not *CYP6M2* are upregulated in RES female abdomens (Unpaired t-tests, *p*<0.00001, *p*=0.0317, *p*>0.05, respectively). Means with SD are shown. (E) Canonical cuticular resistance transcripts *CYP4G16* and *CYP4G17* are not upregulated in the RES colony (Unpaired t-tests, *p*>0.05). Means and SD are shown. *n* represents total number of mosquitoes.

Overall, these data demonstrate that exposure to permethrin resulted in selection for at least metabolic (CYP enzymes) and target site (*kdr* mutations) resistance mechanisms, although it is possible that additional yet unknown mechanisms are at play in RES.

### Males but not blood fed females show fitness costs of insecticide resistance

We next evaluated whether the RES line carried fitness costs in comparison with SUS. We saw no difference in fertility, fecundity or longevity of blood fed females, but observed a modest decrease in longevity of RES females (median lifespan of 20 days) compared to SUS females (median lifespan of 21 days) that were fed only on sugar **(Supplementary Figure 2A-D)**. Similarly, we observed a significant decrease in male longevity where RES males lived a median of 27 days compared to 31 days for SUS males **(Supplementary Figure 2E)**. Together these data suggest that without the additional nutrition from a blood meal, mosquitoes may suffer costs from harboring insecticide resistance traits.

The most substantial difference in fitness was on male mating competitiveness. When SUS and RES males were placed in competition for either RES or SUS females (alternated between experiments to avoid possible biases), RES males were captured *in copula* markedly less often than SUS males, suggesting reduced mating capacity in males that carry insecticide resistance traits **(Supplementary Figure 2F)**. This difference in competitiveness, which was not caused by confounding influences of male body size **(Supplementary Figure 2G)** or assortative mating (as SUS males were more successful regardless of which females were used), may account for why loss of insecticide resistance occurred in the SUS colony.

### RES females are more supportive of *P. falciparum* infection compared to SUS

We next investigated whether RES and SUS females differ in their capacity to support *P. falciparum* infections. Surprisingly, we detected significantly higher numbers of oocysts in the midgut of RES compared to SUS mosquitoes (median 1.9-fold increase) **(Figure 3A)**. Body size was not an important variable in these results as wing length did not differ between the two groups **(Supplementary Figure 3)**. We also measured oocyst size, a good proxy of *Plasmodium* growth rates ^42^, and showed that RES females had 20% larger oocysts on average compared to SUS counterparts 7 days post blood meal (**Figure 3B**). The difference in oocyst numbers and growth was reflected at the level of sporozoites in the salivary glands, where we saw a striking 10-fold increase in the median intensity of infection as well as higher prevalence in RES females compared to SUS 15 days after infection **(Figure 3C)**. These results demonstrate that, even in lines originating from the same genetic background, selection for insecticide resistance can result in increased capacity to host and transmit *P. falciparum*. No differences in longevity were observed between the two infected groups **(Supplementary Figure 3B)**.

**Figure 3:**
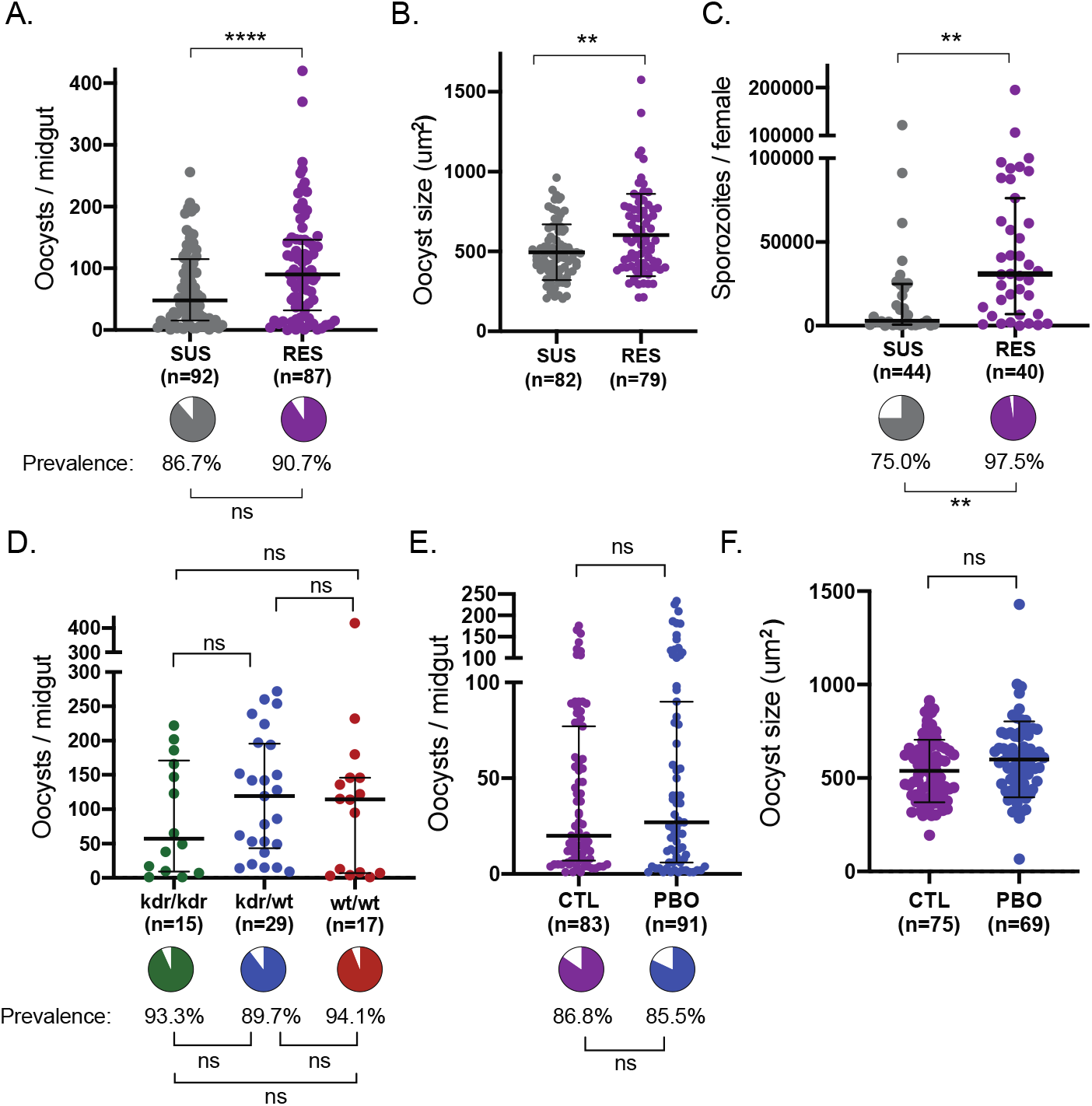
Permethrin resistant females show increased infection intensity and growth rate compared to susceptible females. (A) RES females have higher numbers of oocysts compared to SUS females. (Standard least squares analysis *p*<0.0001). Median and interquartile ranges are shown. Prevalence of infection was not different (pie charts, Nominal logistic *p*>0.05). (B) RES females have larger oocysts compared to SUS females at day 7 post infection (Standard least squares analysis, *p*=0.0031). Mean and SD are shown. (C) RES females have more sporozoites compared to SUS females at day 15 post infection (Standard least squares analysis, *p*=0.0022) and also higher prevalence of sporozoites (pie charts, Nominal logistic, *p*=0.0015). Median and interquartile ranges are shown. (D) *kdr* genotype does not affect either oocyst prevalence (pie charts, Nominal logistic, *p*>0.05), or intensity (Standard least squares analysis with false discovery correction, *p*>0.05). Median and interquartile range is shown. *n* represents total number of mosquitoes. (E) Exposure to PBO prior to an infectious blood meal does not impact intensity (Standard least squares analysis, *p*>0.05) or prevalence (pie charts, Nominal logistic *p*>0.05) of oocysts in RES females. Medians and interquartile ranges are shown. (F) Oocyst size in RES females is unaffected by PBO treatment (Standard least squares analysis, *p*>0.05). *n* indicates number of mosquitoes.

Mimicking lower infection intensities, which are more often observed in field settings, we partially inactivated parasites with heat treatment prior to offering mosquitoes an infectious blood meal. We observed that also under these conditions, RES mosquitoes showed increased prevalence of both oocysts and sporozoites **(Supplementary Figure 2C, D)**, again independent of body size **(Supplementary Figure 2E)**. Due to the low parasite numbers we did not measure oocyst size in these conditions.

### Target site and metabolic resistance do not affect infection in RES females

To address whether the increase in infectious burden in RES was directly related to insecticide resistance mechanisms, we first measured *kdr* frequencies among infected females. We found no difference in oocyst intensity between individuals that were either wild-type, homozygous, or heterozygous for the *kdrL1014F* mutation, suggesting *kdr* is not driving the increased parasite burden in RES **(Figure 3D)**.

We were primarily interested in investigating metabolic resistance mechanisms, hypothesizing that these could either shift investment from processes such as immunity toward the detoxification of insecticide, indirectly favoring parasite development, or cause dysregulation in ROS production in the midgut and therefore have a direct impact on parasite survival. However, when we assessed the role of metabolic resistance by pre-treating RES females with PBO 1h prior to an infectious bloodmeal, we saw no significant impact on oocyst numbers **(Figure 3E**) or size **(Figure 3F)**. Importantly, these results show that PBO treatment in mosquitoes does not favor *P. falciparum* development, which is reassuring for the deployment of this synergist in LLINs ^49^.

Overall, these data suggest that selection for resistance, rather than dominant mechanisms of resistance, favored development of *P. falciparum* parasites, although we cannot exclude that the effects on the parasite are linked to yet unknown resistance mechanisms not detected in our study.

### *Lipophorin* is upregulated in resistant females and linked to increased infection intensity

Although two of the canonical markers of cuticular resistance were not upregulated in RES mosquitoes, we did see a >2-fold increase in *lipophorin (Lp)* transcripts **(Figure 4A)** which were additionally albeit modestly induced 2h after exposure to 2.5X permethrin **(Figure 4B)**. Lp is a lipid transporter whose depletion decreases oocyst numbers and parasite infectivity ^42,43,50,51^. Among its many roles, Lp is thought to transport CHCs in the haemolymph, and therefore could regulate thickening of the cuticle and insecticide penetrance ^52^, so that its upregulation could be partially responsible for both the insecticide resistant phenotype as well as the increase in parasite numbers in RES ^42,43^. A role for Lp in insecticide resistance has, however, never been demonstrated.

**Figure 4:**
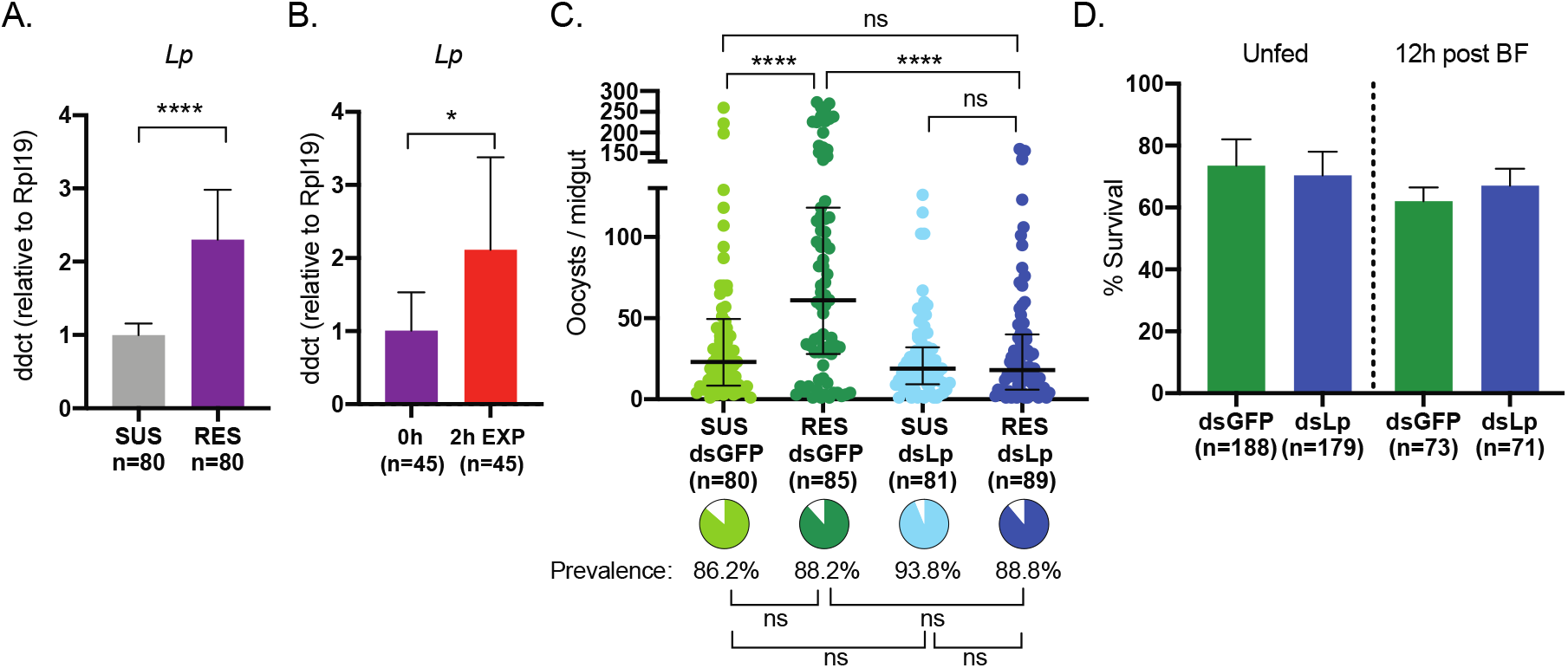
Upregulated *Lp* in resistant females accounts for increased *P. falciparum* infection intensity. (A) Expression of *Lp* is increased in RES females compared to SUS females (Unpaired t-test, *p*<0.0001) and (B) is also increased 2h after permethrin exposure (Unpaired t-test, *p*<0.0275). Mean with SDs are shown. (C) RES controls show higher oocyst intensity than SUS controls, and this difference is eliminated by *Lp* silencing (Standard least squares analysis with false discovery rate corrections, *p*<0.0001, *p*>0.05, respectively). Oocyst prevalence is unaffected by treatment (pie charts, Nominal logistic with false discovery rate correction, *p*>0.05). Medians and interquartile ranges are shown. (D) Knocking down *Lp* in RES does not affect survival to a 2.5X dose of permethrin in either unfed females or 12h after blood-feeding (Chi-squared tests, *p*>0.05). Mean with SD is shown. *n* represents total number of mosquitoes.

We performed RNAi silencing of *Lp* in both RES and SUS females prior to an infectious bloodmeal with the expectation that if increased *Lp* levels are linked to higher infection intensities in RES, we should see that RES and SUS females have the same number of oocysts after *Lp* knockdown. Indeed, while in controls we detected the expected increase in oocyst intensity in RES compared to SUS, *Lp* silencing removed this difference by strongly decreasing the number of oocysts in RES while having more marginal effects in SUS **(Figure 4C)**. Therefore, an excess of Lp is likely to play a role in the increased oocyst numbers observed in RES.

We next determined whether Lp plays a direct role in insecticide resistance. Despite a strong silencing effect (**Supplementary Figure 4A)**, we did not observe any impact on mortality after a 2.5X dose of permethrin in RES females, either unfed or 12h post-blood-feeding **(Figure 4D)**. Reasoning that *Lp* knockdown may have effects on CHC turnover that take longer to impact the cuticle, we also investigated survival upon permethrin exposure 8 days post-injection and after a 5X dose, but still observed no difference in mortality **(Supplementary Figure 4B)**. With the caveat that we did not measure CHC levels in these experiments, these data do not support a role for Lp and CHC transport in the resistance mechanism observed in RES, consistent with our results on the expression of the canonical markers of cuticular resistance **(Figure 2E)**.

### Sublethal permethrin exposure changes lipid profile but not infection outcomes

Given Lp does not play a direct role in survival of RES females after insecticide exposure but is upregulated by permethrin challenge, we hypothesized that this transporter could be involved in a compensatory or recovery response. It is known that permethrin can cause damaging lipid peroxidation ^33^, and transcripts related to lipid biosynthesis and metabolism are upregulated around 72h post exposure ^53^. Lp may mobilize lipids from the fat body following insecticide challenge to provide energy for detoxification and/or to mitigate damage in mosquito tissues.

Based on this reasoning, we performed LC-MS in female fat bodies to investigate differences between RES and SUS females, and in RES females before and after exposure to permethrin. We identified >650 lipids from 6 lipid classes but saw few differences in lipids at either an individual or class level between RES and SUS female fat bodies, indicating that fat stores are similar between these two lines **(Figure 5A, Supplementary Figure 5A)**. When we analyzed RES 24h after a 2.5X exposure to permethrin, however, we saw a decrease in lipids at an individual level **(Figure 5B**) and detected a striking 55% decrease in total lipid abundance **(Figure 5C)**. Strong decreases were also observed in sphingolipids and neutral lipids **(Figure 5D)**, and in several lipid subclasses, including triglycerides which showed a 69% decrease after exposure **(Table 1)**, driving much of the change in total lipids as they account for more than half of lipid abundance in controls.

**Table 1:** Comparisons of lipid subclasses between RES and SUS (unexposed) and RES before (0h) and after permethrin exposure (24h).

| Subclass | # lipids identified |  |  | Mean peak intensity |  |  | FC |  | p value |  |
| --- | --- | --- | --- | --- | --- | --- | --- | --- | --- | --- |
|  | SUS | RES | RES | SUS 0h | RES 0h | RES 24h | RES 0h/SUS | RES 0h /RES | RES 0h/SUS | RES 0h/RES |
| Acyl Carnitine | 7 | 7 | 4 | 7.67E+07 | 1.07E+08 | 6.93E+06 | 1.398 | 15.475 | 0.453 | 0.001 |
| Fatty acid | 1 | 1 | 1 | 2.56E+08 | 2.84E+08 | 3.02E+08 | 1.109 | 0.940 | 0.900 | 0.447 |
| N-Acylethanolamine | 1 | 1 | 1 | 5.23E+06 | 1.56E+06 | 1.19E+08 | 0.298 | 0.013 | 0.520 | 0.484 |
| Phosphatidylethanol | 9 | 9 | 9 | 3.93E+08 | 3.23E+08 | 4.43E+08 | 0.822 | 0.728 | 0.525 | 0.397 |
| Phosphatidylmethanol | 3 | 3 | 3 | 7.09E+07 | 7.35E+07 | 5.55E+07 | 1.037 | 1.325 | 0.904 | 0.464 |
| Wax esters | 2 | 2 | 0 | 4.83E+06 | 2.38E+07 | 0.00E+00 | 4.934 | NA | 0.167 | 0.085 |
| Bis(monooleoylglycerol)phosphate | 7 | 7 | 7 | 5.01E+08 | 5.42E+08 | 4.52E+08 | 1.080 | 1.198 | 0.781 | 0.543 |
| Digalactosyldiacylglycerol | 1 | 5 | 3 | 2.75E+04 | 8.04E+05 | 1.84E+06 | 29.203 | 0.437 | 0.291 | 0.607 |
| Monogalactosyldiacylglycerol | 13 | 13 | 13 | 1.18E+09 | 1.21E+09 | 1.12E+09 | 1.025 | 1.084 | 0.910 | 0.759 |
| Cholesterol Ester | 2 | 2 | 2 | 2.05E+09 | 2.49E+09 | 1.49E+09 | 1.215 | 1.670 | 0.307 | 0.104 |
| Diglyceride | 19 | 19 | 19 | 4.71E+09 | 2.04E+10 | 1.36E+10 | 4.331 | 1.498 | 0.619 | 0.173 |
| Monoglyceride | 3 | 3 | 3 | 1.60E+08 | 6.24E+07 | 1.38E+08 | 0.390 | 0.453 | 0.107 | 0.592 |
| Triglyceride | 106 | 96 | 96 | 8.25E+11 | 6.19E+11 | 1.94E+11 | 0.750 | 3.192 | 0.517 | 0.022 |
| Zymosterol Ester | 2 | 2 | 2 | 4.97E+09 | 3.87E+09 | 1.35E+09 | 0.779 | 2.869 | 0.353 | 0.001 |
| Cardiolipin | 102 | 103 | 103 | 4.14E+10 | 4.82E+10 | 3.28E+10 | 1.165 | 1.471 | 0.642 | 0.293 |
| Lysophosphatidic acid | 1 | 0 | 1 | 1.12E+07 | 0.00E+00 | 1.44E+06 | 0.000 | 0.000 | 0.207 | 0.046 |
| Lysophosphatidylcholine | 8 | 8 | 6 | 1.87E+09 | 1.74E+09 | 7.19E+07 | 0.929 | 24.166 | 0.052 | 0.003 |
| Lysophosphatidylethanolamine | 19 | 18 | 18 | 7.90E+09 | 7.37E+09 | 1.03E+09 | 0.933 | 7.193 | 0.205 | 0.013 |
| Lysophosphatidylglycerol | 3 | 3 | 3 | 2.38E+07 | 2.53E+07 | 1.16E+07 | 1.060 | 2.173 | 0.325 | 0.117 |
| Phosphatidic acid | 9 | 9 | 9 | 2.74E+08 | 2.71E+08 | 1.68E+08 | 0.986 | 1.614 | 0.794 | 0.208 |
| Phosphatidylcholine | 99 | 99 | 97 | 1.38E+11 | 1.76E+11 | 1.18E+11 | 1.275 | 1.492 | 0.382 | 0.153 |
| Phosphatidylethanolamine | 91 | 92 | 93 | 1.09E+11 | 1.15E+11 | 7.96E+10 | 1.056 | 1.445 | 0.318 | 0.202 |
| Phosphatidylglycerol | 25 | 25 | 25 | 1.64E+09 | 1.69E+09 | 1.14E+09 | 1.027 | 1.480 | 0.452 | 0.245 |
| Phosphatidylinositol | 31 | 31 | 31 | 1.01E+09 | 9.85E+08 | 4.06E+08 | 0.974 | 2.429 | 0.600 | 0.046 |
| Phosphatidylserine | 62 | 62 | 62 | 5.26E+09 | 4.78E+09 | 5.90E+09 | 0.909 | 0.811 | 0.584 | 0.544 |
| Ceramide phosphoethanolamines | 1 | 1 | 1 | 4.49E+07 | 6.47E+07 | 1.39E+07 | 1.442 | 4.637 | 0.301 | 0.813 |
| Ceramides | 15 | 15 | 15 | 1.05E+09 | 1.04E+09 | 5.41E+08 | 0.985 | 1.918 | 0.957 | 0.074 |
| Sphingomyelin | 11 | 11 | 10 | 1.78E+09 | 1.37E+09 | 3.98E+08 | 0.767 | 3.431 | 0.456 | 0.004 |
| Sphingomyelin(phytosphingosine) | 4 | 4 | 4 | 1.88E+09 | 1.77E+09 | 1.24E+09 | 0.944 | 1.433 | 0.842 | 0.271 |
| Sphingosine | 1 | 1 | 1 | 4.06E+05 | 3.32E+06 | 4.53E+06 | 8.180 | 0.734 | 0.014 | 0.516 |
| **Total** | **658** | **652** | **642** | **1.15E+12** | **1.01E+12** | **4.54E+11** | **0.877** | **2.221** | **0.716** | **0.0245** |

**Figure 5:**
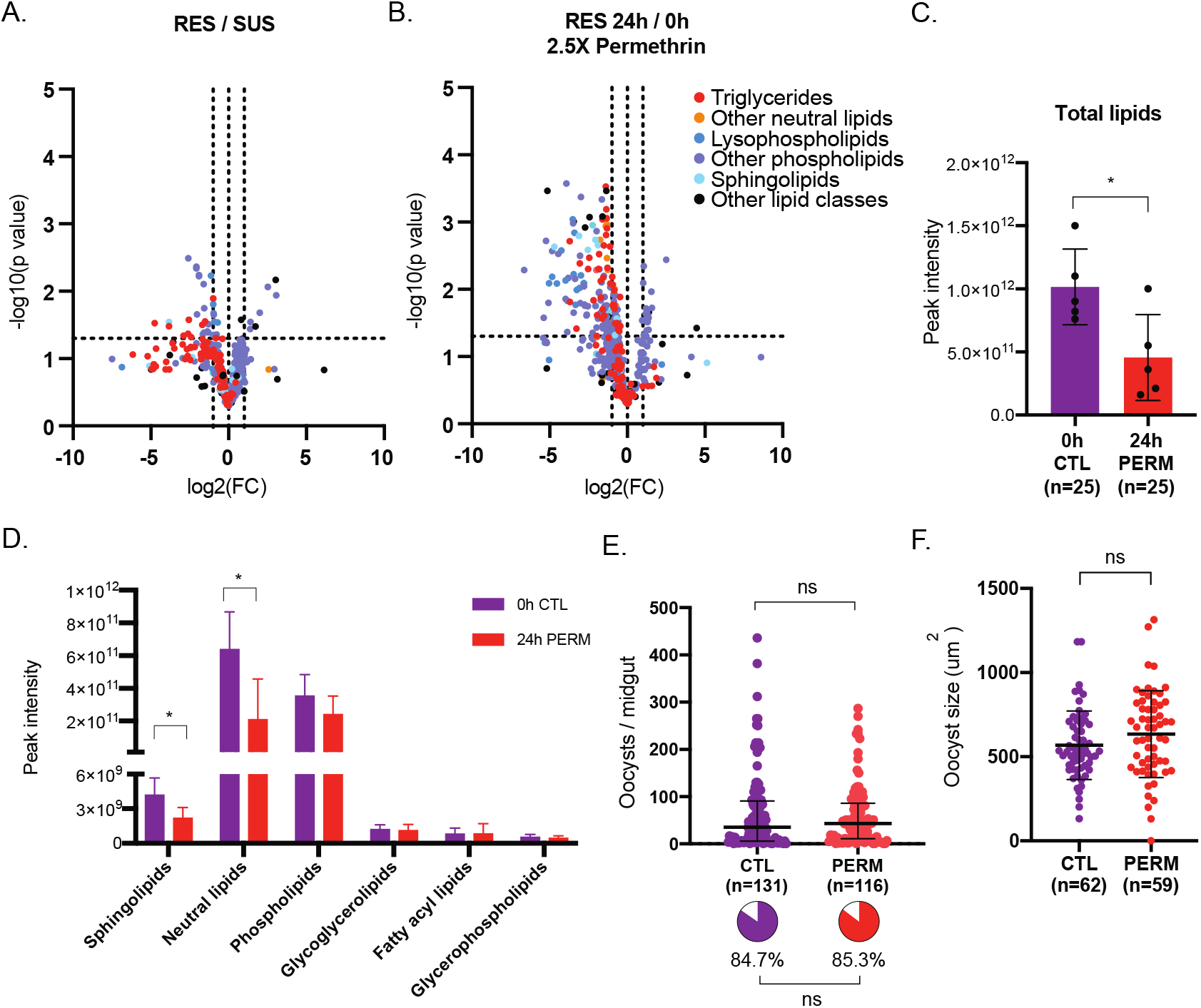
Sublethal permethrin exposure impacts RES lipid profile but not *Plasmodium* infection. (A) Modest significant differences in lipids between RES and SUS female fat bodies were identified. Individual lipids are portrayed according to their fold change (FC) in RES relative to SUS in non-blood fed females, and the statistical significance of the differences in FC (Unpaired t-tests). (B) Several lipid species from a variety of lipid classes and subclasses are decreased in RES female fat bodies 24h after exposure to 2.5X permethrin for 1h. Individual lipids are portrayed according to their FC in RES 24h after exposure relative to unexposed (0h) RES controls, and the statistical significance of the differences in FC (Unpaired t-tests). (C) RES females show decreased total lipids 24h after exposure to 2.5X permethrin for 1h relative to unexposed controls (0h). (One-way ANOVA, *p*=0.0224 and *p*=0.0034, respectively, *p*>0.05 for all other comparisons). Means and SD are shown. (D) 24h following exposure to a 2.5X dose of permethrin, RES female fat bodies showed a significant decrease in abundance of sphingolipids (*p*=0.0327), and neutral lipids (*p*=0.0226) relative to unexposed females. Unpaired t-tests, *p*>0.05 for all other comparisons. Mean with SD is shown. (E) Permethrin exposure 24h prior to an infectious blood meal does not impact oocyst intensity (Standard least squares analysis, *p*>0.05) or prevalence (pie charts, Nominal logistic, *p*>0.05) in RES females. Medians and interquartile ranges are shown. (F) Oocyst size is also not impacted by permethrin exposure 24h prior to infection (Unpaired t-test, p>0.05). Means and SD are shown. *n* represents total number of mosquitoes.

As there are known links between the availability of lipids and *P. falciparum* infections ^42-44^, we next wanted to assess the impact of permethrin challenge on parasites. Interestingly, we observed no effects on parasite numbers, size or prevalence in RES females that were exposed to 1X permethrin either 24h or 5h prior to infection **(Figure 5E, F, Supplementary Figure 5B, C)**, suggesting that remobilized lipids are not available to the parasite for either growth or survival. The lack of an effect of permethrin exposure on infection is also perhaps a further indication that metabolic resistance mechanisms, even when further activated by direct insecticide exposure ^53-56^, do not influence *Plasmodium* survival and growth in the mosquito vector.

## Discussion

This study demonstrates that selection for insecticide resistance can result in increased *P. falciparum* numbers and growth rate. RES and SUS colonies were derived from the same genetic starting material and were strikingly similar at the level of life history where we identified very few differences in fitness between them. Consistently, these lines showed strong similarities also in terms of their lipidomic profiles, supporting the concept of this experimental design which aimed to create genetically similar colonies differing mainly in their insecticide exposure and response. Permethrin resistance was lost from G3VK when it was not actively selected for, and we speculate that this is likely due to the significantly reduced competitiveness of RES males during mating. Although mating success has been positively correlated with insecticide resistance previously, this is attributed to cuticular resistance and particularly CHC abundance ^57^, which does not appear to contribute to the resistance phenotype in our study. Other observed fitness costs, such as decreased longevity of sugar-fed males and females, were unlikely to affect fitness under our rearing conditions. There may however be other discrete fitness costs we have not investigated (potentially during larval development) that also play a role in the loss of resistance over time.

*P. falciparum* infection outcomes were not affected by *kdr*, a result which did not surprise us given the specific mechanism of target site resistance is unlikely to directly affect parasite development in the midgut. Although some other studies have seen increased *Plasmodium* infection rates or intensity when at least one *kdr* allele was present ^24,58^, others have identified no association^26,59^, Further, one investigation demonstrated no impact of functional knockdown of *para* on infection, instead suggesting that the *kdr* mutation could be in genetic linkage with a gene that impacts immunity ^25^. Given our results, such linkage is however unlikely to have occurred in our selection process.

We were surprised that neither PBO nor permethrin exposure impacted the establishment of *P. falciparum* in the midgut, as both treatments can affect the redox state in the mosquito and ROS are known to limit *Plasmodium* infections ^35,60^. Permethrin exposure leads to ROS production ^33,61^ and induction of CYPs. CYP activity (inhibited by PBO) can increase superoxide radicals and hydrogen peroxide levels through oxygenation reactions ^32^. It is possible that these treatments affect ROS in other tissues but not the midgut, and therefore may not impact parasite development. Regardless, it is reassuring that PBO exposure did not exacerbate *P. falciparum* infection, as it is beginning to be widely used as a synergist in LLINs ^49^.

Increased infection intensities were instead associated with *Lp* upregulation in the resistant population, as knocking down this lipid transporter eliminated differences in oocyst numbers between RES and SUS. Lp has been previously demonstrated to affect parasite survival in the midgut ^42^, but the mechanisms by which this effect is carried out are yet unknown. We could not establish a direct link between Lp and insecticide resistance per se as Lp depletion did not restore mortality in RES, but Lp may still play a compensatory role in tolerating insecticide pressure. Specifically, Lp may be involved in lipid mobilization after pyrethroid exposure, as we observed a decrease in fat body lipid stores 24h after permethrin treatment. Mosquitoes may deploy lipids from the fat body to other tissues to aid or accelerate their recovery from insecticide challenge, possibly repairing damage caused by lipid peroxidation ^33,34^. Such effects may not be essential for survival after acute exposure, explaining why Lp did not affect mortality, but if lipid mobilization by Lp ultimately impacts fitness in females exposed to insecticides, this may explain its upregulation in RES after permethrin challenge. Such a compensatory mechanism could have a more prominent role in field settings where mosquitoes are subject to multiple stressors. As an additional or alternative hypothesis, it is possible that upregulated *Lp* expression in RES is genetically linked to other resistance mechanisms that we did not identify in our analysis, explaining the increased parasite infections in resistant females.

We expected that the changes we observed in the lipid profile after 1X permethrin exposure might impact *Plasmodium* infections, but we saw no difference in oocyst intensity or size in controls compared to permethrin-exposed females (**Figure 5E, F**). This may be because exposure occurred close to the blood meal, which contributes additional lipid resources to the mosquito, possibly masking permethrin’s impact. It would be interesting to explore the effect of permethrin exposure throughout parasite development, and to use a higher permethrin dose such as the 2.5X dose that was seen to stimulate alterations in the lipid profile of RES females (**Figure 5B-D**). As previously mentioned, other studies have demonstrated that sublethal pyrethroid exposure can decrease *Plasmodium* intensities ^29-31^. Differences in the timing of insecticide exposure prior to blood-feeding, or in the mechanisms of insecticide resistance in different mosquito strains could underlie these discrepancies.

Given the multiple resistance mechanisms observed in these mosquitoes derived from the same starting field-derived, highly resistant population of *An. coluzzii*, our study takes into account (but does not directly address) possible synergy between these mechanisms that may influence mosquito physiology, and is more controlled than comparisons of resistant and susceptible lines from different geographic and temporal origins. Additional work is needed to determine how commonly *Plasmodium* infection outcomes are influenced by selection for (or loss of) insecticide resistance in order to provide more detailed information on malaria transmission dynamics in field-relevant conditions. Despite this caveat, our data demonstrate that insecticide pressure can at least in some circumstances lead to increased vector competence for *P. falciparum*. While this should not be overinterpreted to suggest that insecticide resistance increases transmission likelihood in natural populations, our findings stress the urgency of insecticide resistance monitoring paralleled by careful consideration of discrete consequences of insecticide resistance selection.

## Materials and methods

### Mosquito lines and rearing

#### Generating resistant and susceptible lines

Blood fed females were collected from inside walls from the Vallée du Kou village 5 (VK5), and allowed to oviposit. Eggs were then mailed to Boston where they were floated. Adults that were reared from these eggs were crossed with mosquitoes from the Mopti colony in 2015. The resulting line was outcrossed again to adults reared from VK5 in 2017 (**Supplementary Figure 1A**). This line was outcrossed once more to VK5-derived mosquitoes in 2018 to yield the VK colony. As the VK colony could never be deselected for insecticide resistance (**Supplementary Figure 1B**), it was crossed with G3 mosquitoes to yield the G3VK line (**Figure 1A, Supplementary Figure 1A**). To select the resistant RES line, G3VK females were exposed to a 1X dose of permethrin between 3-7 days old over multiple generations. Surviving females were blood fed and contributed to the next generation. The susceptible SUS line was reared in concert with the RES line, but was never exposed to permethrin.

#### Rearing conditions

Mosquitoes were maintained in a 27°C incubator with 70-80% humidity and a 12h light: 12h dark cycle. Adults were given 10% glucose and water ad libitum and fed on human blood (either from a human volunteer or from Research Blood Components, Boston, MA). Larvae were fed a mixture of Tetramin fish flakes and pellets.

#### Insecticide resistance bioassays

For selections, mosquitoes were exposed to a 1X dose of permethrin using the WHO bioassay guidelines^62^ with 0.75% (1X) permethrin, 1.875% (2.5X) or 3.75% (5X) permethrin-impregnated papers for one hour, unless otherwise noted. Females were typically 2-5 days old for exposure. For synergist assays, Grade 1 Whatman papers were impregnated with either 4% PBO or 8% DEM in acetone. 4-5 day old females were pre-exposed to PBO or DEM for 1 hour immediately prior to permethrin exposure at either a 1X, 2.5X or 5X dose. Mortality was evaluated 24h after exposure. For infection experiments, females were exposed to PBO for 1h and then were offered an infectious blood meal within one hour post-exposure. For ds*Lp* experiments in **Figure 4D**, unfed females were exposed at 5 days old (4 days post injection) and blood fed females were offered blood at 5 days old (4 days after injection) and exposed the subsequent morning 12h later (5 days post injection).

#### Egg development assays

Virgin females were given access to a blood meal and their ovaries were dissected 3 days later. Eggs were counted using a Leica M80 dissecting microscope.

#### Fertility assays

To ensure females were mated, they were caught *in copula* during natural matings with males from the same colony. They were then offered a blood meal after which unfed females were removed. Females were then given ad libitum access to 10% glucose solution and water for 2 days prior to allowing oviposition in individual cups lined with filter paper. Once laid, eggs were stimulated daily by spraying water and allowed to hatch for a minimum of 4 days. Fertility was assessed by counting and scoring eggs under a Leica M80 dissecting microscope, and additionally noting the presence or absence of hatched larvae. If any female had no fertile embryos, we verified her mating status by checking microscopically for the presence of sperm in the spermatheca.

#### Longevity assays

Pupae were sexed prior to being placed in a cage of 50-100 individuals. Mortality was assessed every 1-2 days. Mosquitoes were given access to 10% glucose solution and water *ad libitum*. For blood fed female longevity assays, virgin females were blood fed once, at 7 days old. Mortality was only assessed following the blood meal.

#### Mating competition assays

Virgin males and females were separated by sex sorting as pupae. Approximately 100 males of each the RES and SUS line were coated in either pink or yellow fluorescent dust (colors swapped between replicates) at 3-4 days old. This was done by anaesthetizing males on ice and gently shaking them in a glass dish coated with fluorescent dust. We found this method resulted in better recovery and swarming compared to the published syringe method ^63^, which in our hands resulted in very limited swarming behavior. Males were then mixed into a single cage for mating competition assays. 5-10 females were released into the cage at a time, and when they were found *in copula*, they were caught and removed from the cage. After swarming and mating for approximately one hour, all couples were evaluated for the color of the male, and whether an autofluorescent mating plug was received by the female using a Leica M80 fluorescence dissecting scope. When no plug was received, the couple was excluded from analysis.

#### P. falciparum cultures and infection

##### Parasite cultures

NF54 *P. falciparum parasites* (confirmed by PCR^64^) were obtained with permission from Carolina Barillas-Mury, National Institutes of Health, MD, USA. Asexual parasites were cultured at 37°C using human erythrocytes at 5% hematocrit (Interstate Blood Bank, Memphis TN) in RPMI 1640 with supplemented 25mM HEPES, 10mg/L hypoxanthine, 0.2% sodium bicarbonate, and 10% heat-inactivated human serum (Interstate Blood Bank) incubated with 5%O2, 5% CO2, and balanced N2 for up to 8 weeks, in accordance with published protocols^65,66^. To stimulate gametocytogenesis, parasitemia was increased above 4% and cultures were maintained for 14 to 20 days such that stage IV and V gametocytes are concentrated.

##### Mosquito infection

Mosquitoes were starved overnight prior to receiving an infectious blood meal. Gametocyte cultures were pipetted into membrane feeders attached to a hot water system to keep the cultures warm. Caged female mosquitoes were allowed to feed on infected blood for up to one hour, after which unfed females were removed. Blood fed females remained inside a custom glove box (Inert Technology, Amesbury, MA) with access to 10% glucose solution until dissection.

##### Oocyst counts and measurements

At 7 days post infection, females were removed from cages by vacuum aspiration into 80% ethanol and frozen to ensure death. Females were then dissected in PBS, midguts were stained with 2mM mercurochrome and imaged at 100X on an Olympus Inverted CKX41 microscope, and oocysts were later counted and measured on FIJI^67^. All measurements within replicates were taken by the same person for consistency.

##### Sporozoite counts

At 15 days post infection, females were removed from cages by vacuum aspiration into cold PBS and carefully decapitated to ensure death. Salivary glands were dissected and homogenized manually with a pestle for 30 seconds, then pelleted by centrifugation for 10 mins at 4C, 8000 x g, and resuspended in 40µL PBS. 10µL was then pipetted onto a disposable haemocytometer. Sporozoites were visualized and counted on an Olympus Inverted CKX41 microscope at 200X with phase-contrast microscopy for each female.

#### Wing length measurement

Wings were imaged and measured from the proximal wing notch to the distal tip of the third cross vein using ImageJ^67,68^. All measurements within each experiment were taken by the same person for consistency.

#### DNA extraction and genotyping

After dissecting midguts for oocyst counts, carcasses were preserved in ethanol until DNA extraction with the Qiagen Dneasy kit. DNA samples were genotyped for the *kdr*L1014F mutation using Taqman probes generously donated by Dr. Hilary Ranson’s laboratory at the Liverpool School or Tropical Medicine.

#### Transcriptional assays

Female abdomens were dissected in pools of 8, except for in Figure 4B where Lp transcripts are analyzed after exposure, where pools of 5 female abdomens were used. Tissues were collected in TRI reagent (Thermo Fisher Scientific) and stored at -80°C. RNA was extracted according to manufacturing instructions with an additional three ethanol washes of pelleted RNA. Following resuspension, RNA was treated with Turbo DNAse (Thermo Fisher Scientific), quantified with a Nanodrop 2000C (Thermo Fisher Scientific), and then 2µg were used in a 100µL cDNA synthesis reaction, following standard protocols. We designed primers for qRT-PCR (QuantStudio 6 pro, Thermo Fisher Scientific) using NCBI PrimerBLAST^69^ and we used primers in **Supplementary Table 1**. Relative quantification was determined using the 2^-(dCt)^ equation, using RpL19 as the standard, except in Figure 4A, B where ddct is used to normalize disparate replicates with variation between controls. All primers were used at 300nM with the exception of 900nM for RpL19 R.

#### dsRNA synthesis

The eGFP PCR fragment was amplified from plasmid pCR2.1-eGFP as described by Baldini et al. 2013 ^70^. Plasmid pLL10-Lp, a gift from Miranda Whitten and Elena A. Levashina (Max Planck Institute for Infection Biology, Berlin), was used as a template to amplify a 600 bp fragment of Lp (AGAP001826) corresponding to bases 9452–10051 cDNA using a primer matching the inverted T7 promoters: 5’–TAATACGACTCACTATAGGG–3’. PCR product size was verified by gel electrophoresis. Using the PCR product as a template, dsRNA was transcribed and purified using the Megascript T7 *in vitro* transcription kit (Thermo Fisher Scientific) as previously described ^71^.

#### Knockdowns with dsRNA

690ng of dsRNA was injected into adult females within 1d of eclosion using a Nanoject II (Drummond), at a concentration of 10ng/nl.

#### Lipidomics

##### Sample preparation

Fat bodies were collected in methanol in pools of 5 from 4-5 day old non-blood fed females from either the SUS colony, or from the RES colony before or 24 hours after exposure to a 2.5X dose of permethrin using the WHO bioassay. Five pools were collected for each group. Briefly, tissues were homogenized in methanol using a bead beater, before transfer to a glass vial and addition of 4mL chloroform. Samples were vortexed for 1 min prior to addition of 2mL ultrapure water, and then vortexed again. Vials were centrifuged for 10 minutes at 3000 x g and the chloroform phase was separated and used for lipidomics. Samples were dried under N2 and resuspended in 60μL chloroform.

##### LCMS

LC–MS analyses were modified from^72^ and were performed on a ThermoFisher QE+ mass spectrometer coupled to an LC (Ultimate 3000, Thermo Fisher). 20μL of sample was injected onto a Biobond C4 column (4.6 × 50 mm, 5 μm, Dikma Technologies) kept at 25°C. The mobile phases for the LC were A (5% methanol, in water with, for positive mode: 5mM ammonium formate and 0.1% formic acid; and for negative mode : 0.03% ammonium hydroxide) and B (5% water, 35% methanol, 60% isopropanol with, for positive mode: 5mM ammonium formate and 0.1% formic acid; and for negative mode : 0.03% ammonium hydroxide). The LC gradient was as follows: Flow rate was set 100 μl min^−1^ for 5 min with 0% mobile phase B (MB), then switched to 400 μl min^−1^ for 50 min, with a linear gradient of MB from 20% to 100%. The column was then washed at 500 μl min^−1^ for 8 min at 100% MB before being re-equilibrated for 7min at 0% MB and 500 μl min^−1^. Ionization in the MS source was done by heated electrospray ionization and the MS was acquiring data in top 5 automatic data dependent MSMS mode. The mass spectrometer measured data in full-MS mode at 70,000 resolution over an *m*/*z* range of 150 to 2000 and MS2 at a resolution of 35,000. Each sample was run twice, once in positive mode and once in negative mode. Lipids were identified and signal integrated using the Lipidsearch © software (version 4.2.27, Mitsui Knowledge Industry, University of Tokyo). Integrations and peak quality were curated manually before exporting and analyzing the data in Microsoft Excel.

#### Statistical analysis

Longevity data were analyzed in GraphPad Prism 8. Oocyst size and numbers, egg numbers, and sporozoite intensity and prevalence data were analysed in JMP Pro 14 with multifactorial models. Zeroes were excluded when analyzing intensity data. Replicate and wing length, where relevant, was incorporated into these models, as well as full factorial interacting factors. When these variables were not significant to the model, they were removed from the analysis through backward elimination.

## Acknowledgements

Thanks to Abdoulaye Diabaté and his group for their longstanding collaboration and for making these studies possible by sending us *An. coluzzii* eggs from VK5 in Burkina Faso for several years. Thanks to Douglas Paton for helpful feedback during the duration of the project. We are grateful to William Robert Shaw for careful reading of the manuscript. Funding for this study was provided by a joint Howard Hughes Medical Institute and Bill and Melinda Gates Foundation (BMGF) Faculty Scholars Award (Grant ID: OPP1158190) and by the National Institutes of Health (NIH) (award numbers: R01 AI104956 and R01 AI124165) to F.C. as well as by the Natural Sciences Engineering Research Council of Canada (Award number PGSD3-488065-2016). F.C. is an HHMI Investigator.

