## Supplementary Information for "Selection for insecticide resistance can promote *Plasmodium falciparum* infection in *Anopheles*"

### Supplementary Figures for Adams et al. 2022

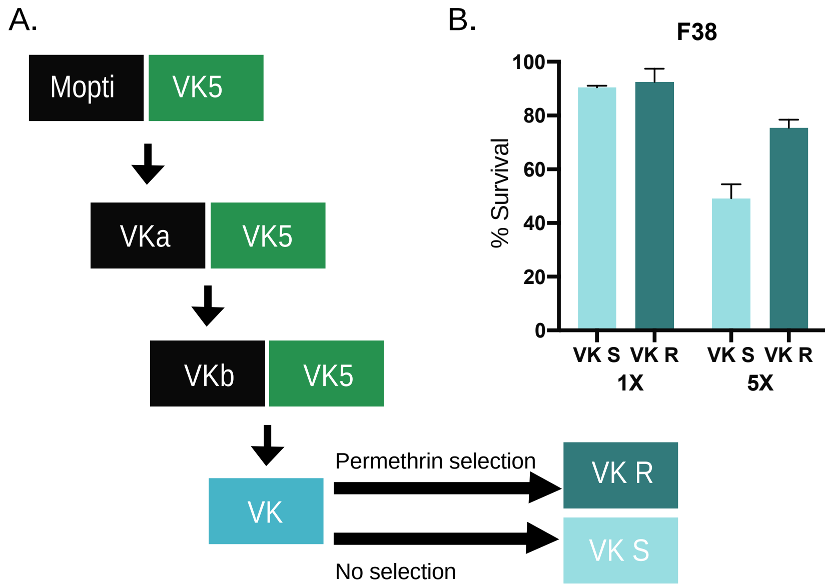


**Supplementary Figure 1:** (A) Scheme of the overall selection process. *An. coluzzii* mosquitoes derived from the VK5 region in Burkina Faso were originally outcrossed to the Mopti strain, and subsequently backcrossed to VK5 mosquitoes twice to establish the VK colony. VK mosquitoes were either exposed to permethrin every generation (VK R) or left in the absence of insecticide pressure (VK S). (B) Despite the lack of insecticide pressure, VK S maintained high levels of insecticide resistance compared to VK R females even after 38 generations, assessed by WHO bioassays using either a 1X or 5X dose of permethrin for 1h. Means and SD are shown.

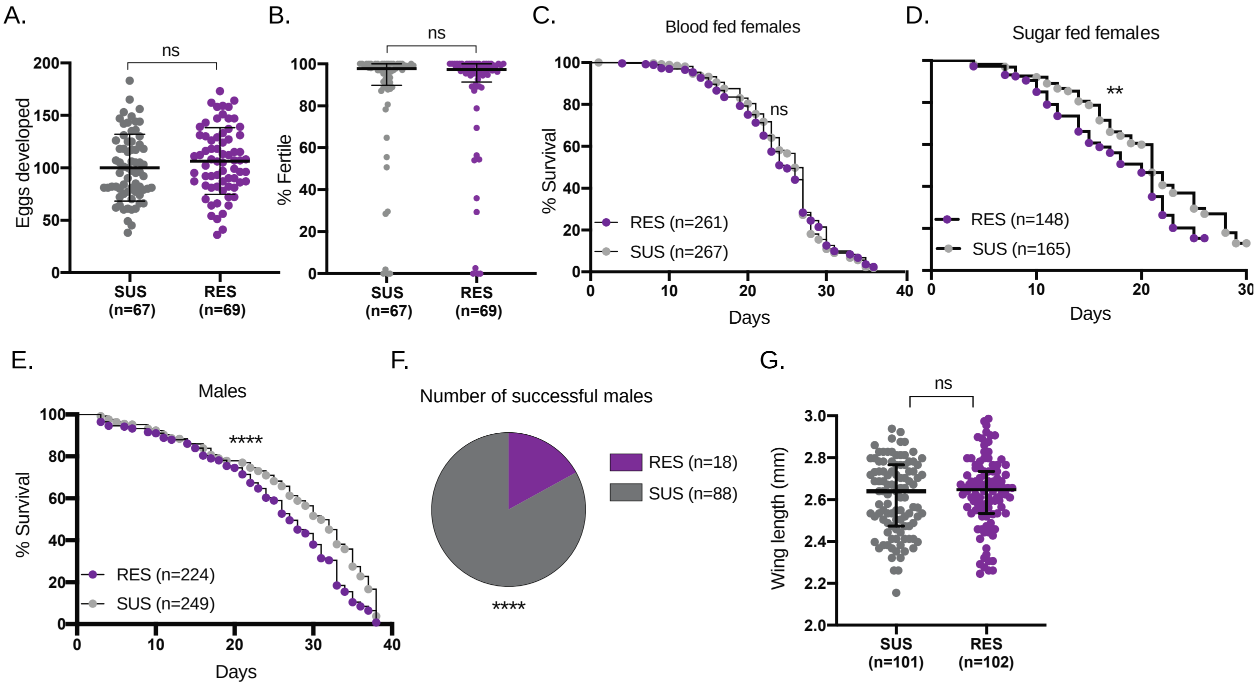


**Supplementary Figure 2: Permethrin-resistant males, but not blood fed females, bear fitness costs.** (A) Number of eggs developed by RES and SUS females are not different (Standard least squares analysis, $p>0.05$). Mean and SD are displayed. (B) Fertility is not different between RES and SUS females (Standard least squares analysis, $p>0.05$). Mean and SD are displayed. (C) Longevity after blood-feeding is not different between RES and SUS females when mortality was recorded after 7 days old were given a blood meal (Log-rank test, $p>0.05$). (D) RES females have decreased longevity compared to SUS females when fed on solely sugar (Log-rank test, $p=0.0016$). (E) RES males have decreased longevity compared to SUS males (Log-rank test, $p<0.0001$). (F) SUS males successfully mate with females more frequently than RES males during mating competition assays (Chi-squared test, $p<0.0001$). (G) Wing length, a good proxy for body size, does not affect male mating competitiveness (Unpaired t-test, $p>0.05$). *n* represents total number of mosquitoes. Mean and SD are shown.

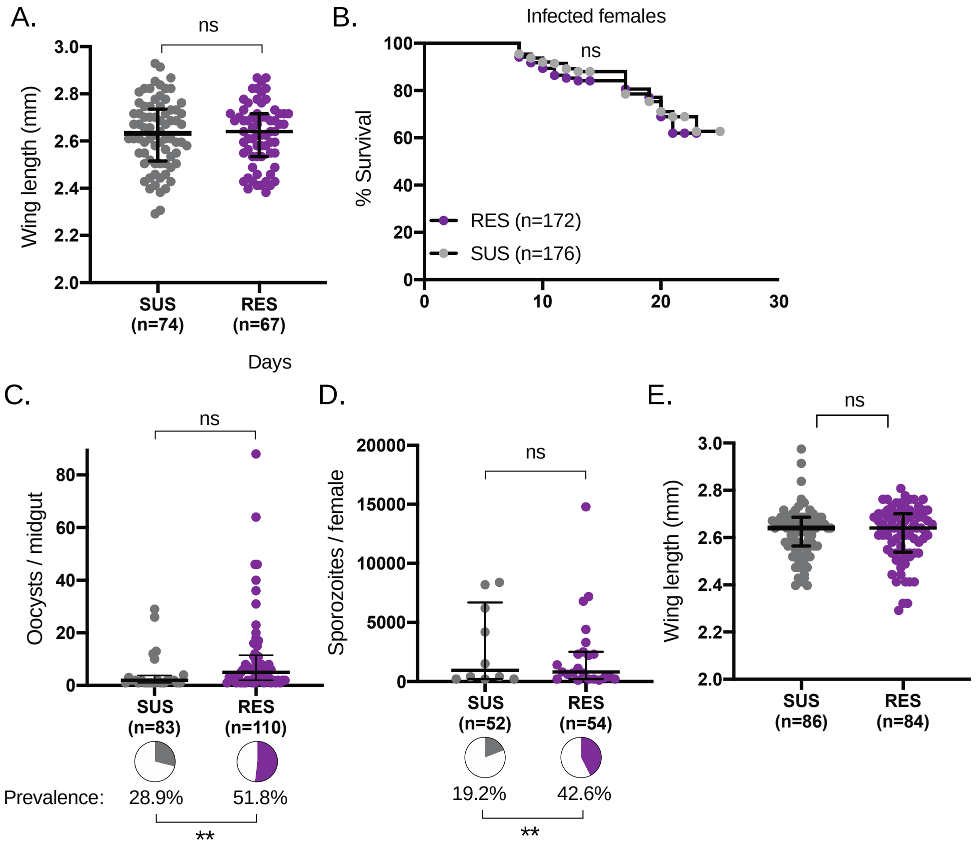


**Supplementary Figure 3:** (A) Wing lengths of RES and SUS females were not different in infection experiments (Unpaired t-test, $p>0.05$). Mean and SD are shown. (B) Longevity of infected females was not different between RES and SUS females used for infection experiments in Figure 3. (Log-rank test, $p>0.05$). Mortality was recorded after an infectious blood meal at 7 days old, and assays were terminated when females were 26 days old due to experimental limitations. (C) When parasites were partially heat-inactivated to approximate field-like lower infection intensities, RES females had increased oocyst prevalence (Nominal logistic, $p=0.0025$). Intensity of infection was instead comparable between the groups (Standard least squares analysis, $p>0.05$). (D) After partial heat-inactivation, RES females had higher prevalence of sporozoites compared to SUS females, (Nominal logistic, $p=0.0041$), though not higher numbers (Standard least squares analysis, $p>0.05$). (E) Wing lengths of RES and SUS females were not different in low intensity infection experiments (Unpaired t-test, $p>0.05$). *n* represents total number of mosquitoes.

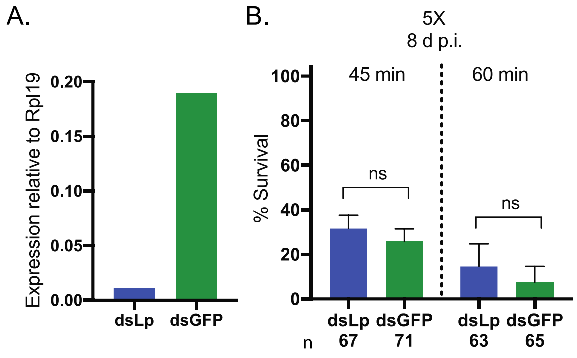


**Supplementary Figure 4:** (A) Expression of *Lp* in ds*Lp* groups is <5% of expression in ds*GFP* control group. (B) No difference in mortality is observed between ds*Lp* and ds*GFP* females exposed to a 5X dose of permethrin 8 days after knockdown, for either 45 minutes or 60 minutes (Chi-squared tests, $p>0.05$). *n* represents total number of mosquitoes.

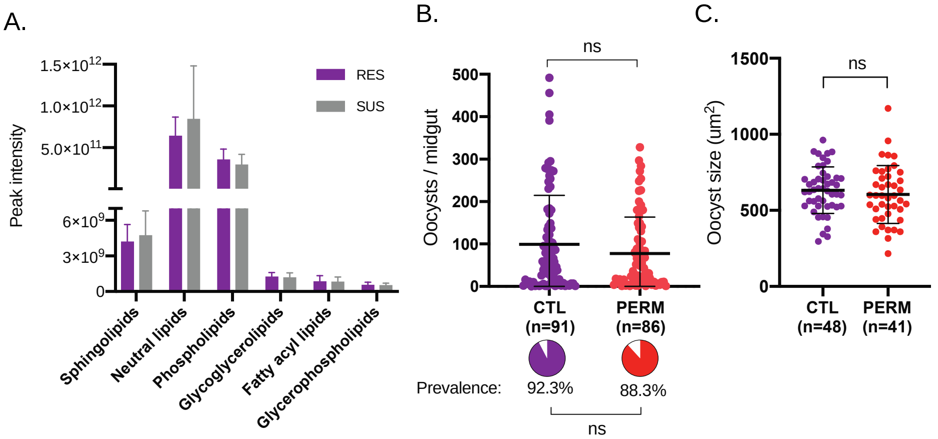


**Supplementary Figure 5:** (A) RES and SUS female fat bodies do not differ in lipid class abundance (Unpaired t-tests, $p>0.05$ for all comparisons). Mean with SD is shown. (B) Exposure to 1X permethrin 5h prior to an infectious blood meal does not impact oocyst intensity (Standard least squares analysis, $p>0.05$) or prevalence (Nominal logistic, $p>0.05$) in RES females. Medians and interquartile ranges are shown. (C) Oocyst size is also not impacted by permethrin exposure 5h prior to infection (Unpaired t-test, p>0.05). *n* represents total number of mosquitoes.

### Supplementary Table 1: Primer sequences for qRT-PCR.

| | | |
|---|---|---|
| *CYP4G17* | F | 5' TGACGGTGGACATTCTGCTC |
| | R | 5' GTCACACATTTTCATGACAGCCA |
| *CYP4G16* | F | 5' GAAGTTGCGTCGGACGTAAATCTA |
| | R | 5' GTCTTCGATTTGCGTTGACGTGGTTC |
| *CYP6M2* | F | 5' TACGATGACAACAAGGGCAAG |
| | R | 5' GCGATCGTGGAAGTACTGG |
| *CYP6P3* | F | 5' TGTGATTGACGAAACCCTTCGGAAG |
| | R | 5' ATAGTCCACAGACGGTACGCGGG |
| *CYP6Z2* | F | 5' CCACGCAATTGCATTGGTCT |
| | R | 5' TTCTACGCGCATGGGGAAAC |
